# Passive muscle forces in *Drosophila* are large but insufficient to support a fly’s weight

**DOI:** 10.1101/2025.04.29.651225

**Authors:** Ninghan Wang, Helene Babski, Jonathan Elliot Perdomo, Sarah Beth McMahan, Arun Ramakrishnan, Tirthabir Biswas, Vikas Bhandawat

## Abstract

Movement of a limb is shaped by active forces generated by muscle contraction but also by passive forces within individual muscles and joints. In small animals such as insects, the contribution of passive forces to limb movement can match the active forces. However, most measurements of passive forces are limited to the femur-tibia joint in large insects. Here we take advantage of genetic tools in *Drosophila* to measure passive torques at multiple joints in the fly’s leg. We genetically inactivate all the motor neurons to assess passive forces. We find that the passive torques are well approximated by a linear spring, i.e., the passive torques linearly increase with angular deviation from the rest angle. The torques are much larger than the gravitational torque due to the leg itself. We estimate that the passive torques are seventy times smaller than necessary to support the weight of the animal. We also inactivated all the motor neurons in a freely standing fly and found that, as predicted from the model, the fly falls when the motor neurons are inactivated. We found that the height at which a fly stands, and, therefore the active forces vary. The fly’s height affects the time to initiate a fall. The time it takes for the fall is consistent with the active forces decaying with a time constant of ∼100 ms. Thus, although passive forces are strong and will have a large effect on limb kinematics, they are not strong enough to support the weight of the fly.

## Introduction

Movement of an animal’s limb is shaped not only by active forces generated by muscle contraction but also by other forces including inertial forces produced by gravity, recoil of one limb segment resulting from the movement of another segment, and passive forces in the muscles and joints [1-11]. The relative contribution of these forces depends on the size of the limb [1]: both inertial forces and active forces scale with the mass of the limb which in turn scales with the volume of the limb and therefore depends on the third power of limb length (L^3^). In contrast, passive forces scale with either the cross-sectional area of the muscle or the surface area of the joint and therefore have a L^2^ dependence. As the size of the limb decreases, the relative contribution of passive forces which is inversely proportional to L (L^2^/L^3^) increases [1, 12]. This scaling law implies that both the movement and posture of low mass limbs are influenced more by passive forces than by inertial forces. Therefore, understanding the control of small biomechanical systems such as the insect leg or human wrist [13-15] or ankle [8, 16] is impossible without first understanding passive forces.

Recent work has begun to explore the consequences of large passive muscle forces in the context of the control of an insect limb. One such consequence is that the insect limb assumes a gravity-independent position because the passive forces are strong enough to counteract the effect of gravitational forces [1]. Another consequence is that active muscle forces are necessary through the entire duration of a leg’s movement: In large limbs such as a human leg, active forces are necessary only at the beginning of movement because inertial forces (or momentum) can carry the leg through the rest of its movement. In insect legs, inertial forces are weak compared to passive muscle forces causing the limb to return to its position determined by passive forces. Therefore, active forces are necessary throughout the movement. Another recent study has shown that passive forces play an important role in stopping the limb accurately [2]. A similarly important role for passive forces was found in the aimed movement of locust legs [3]. Efforts to measure passive forces in insect legs have been primarily limited to the femur-tibia (FTi) joint of large insects [4-7]. These efforts have found that the passive forces at the FTi joint arise from both the elastic properties of the cuticle and from the passive forces within the muscle itself. A limitation to our current understanding of passive muscle forces is a lack of similar measurement at other leg joints. Another limitation is that much of the effort in the measurement of passive forces has been limited to large insects; if scaling law holds true, the effect of passive forces is likely to be even more consequential in small insects such as *Drosophila* which are several orders of magnitude lighter than larger insects. In particular, the passive forces measured in the FTi joint of multiple arthropods suggest that the passive forces might be sufficient to support the weight of the respective animals [4]. If scaling laws hold, the passive forces in the *Drosophila* leg are even more likely to support the weight of a fly.

In this study we measure passive torques in the leg of an intact *Drosophila* by reversibly inactivating all motor neurons and estimating passive torques at each joint from the resulting leg configuration. We show that the torques at each joint can be modeled as a linear spring. The passive torques cannot support the weight of the fly; torques at least 70 times larger are necessary. This prediction was verified by silencing all the motor neurons in a freely walking or standing fly. Unexpectedly, the same fly stands at vastly different heights. Consistent with the idea that the active forces necessary to maintain a given height increases with the height of the fly, the time it takes to initiate a fall increases with the height of the fly. The rate at which a fly falls is consistent with the active forces decaying with a time constant of ∼100 ms Apart from reporting the passive forces in an important model system, our methodology is novel and complements the ones employed in larger insects because we were able to determine the torques at each joint in a completely intact leg. Moreover, employing genetic methods allows an assessment of the role of passive forces not possible otherwise.

## Results

We reasoned that the rest position of the leg when a fly is attached to a tether and suspended in air would be determined by the equilibrium between torques due to active muscle forces, gravity, and passive forces. When all the motor neurons are genetically inactivated, the only two forces remaining are the gravitational force and the passive forces; therefore, the equilibrium angle for any joint is determined by the torques due to these forces. We manipulated the gravitational torque by rotating the body; in response, the joint angle at rest changed. By calculating the gravitational torque at different rest positions, we can estimate how passive torque varies with joint angle.

The flies of the genotype *VGlut-Gal4; UAS-GtACR1* express the light-activated anion conducting channel - *Guillardia theta* anion-conducting receptor1 (GtACR1)[17] in all glutamatergic neurons. Since the motor neurons in *Drosophila* are glutamatergic, they are silenced when the fly is illuminated with green light. We rotated a tethered fly of the above genotype using a precision rotational drive while recording its leg position using two cameras (Figure 1A). Inactivation causes a change in the leg’s rest position; however, in preliminary experiments, the body rotation did not have a large effect on the rest positions of the leg following inactivation. This result is consistent with the one already reported for stick insects and shows that passive forces within the leg are much larger than the gravitational force on a leg and dominate limb position [1]. Therefore, in this scenario, changing the fly’s orientation does not affect the leg rest position because changing the fly’s orientation will only affect gravitational forces which are too weak compared to the passive forces.

**Figure 1.**
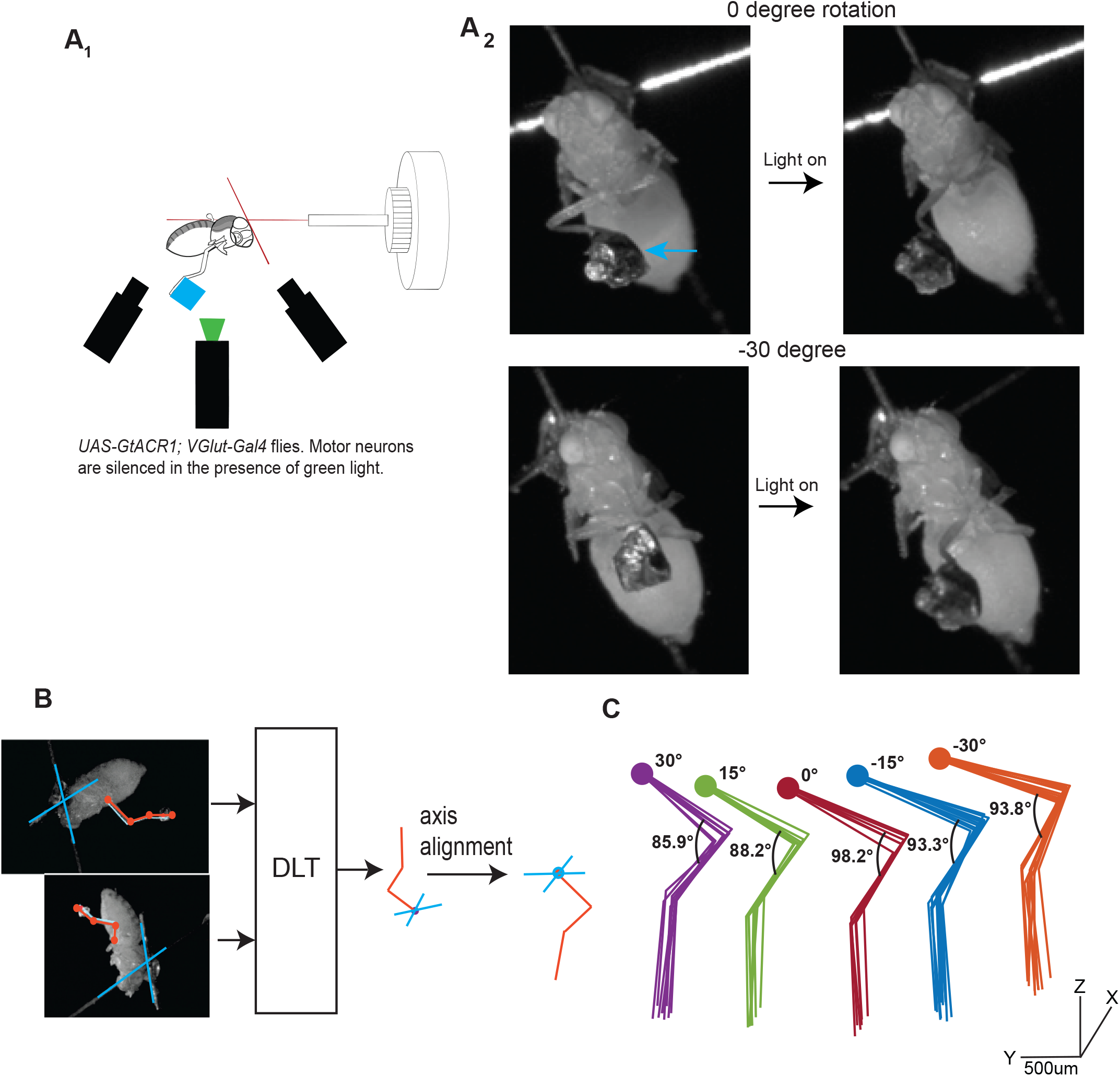
Passive forces in the fly leg are much stronger than gravitational force. **A**_**1**_.Experimental set-up. *UAS-GtACR1; VGlut-Gal4* flies expressing light-activated anionic channels (*GtACR1*) under the control of *VGlut* promoter are rotated. Motor neurons in these flies are inactivated when stimulated with green light. One leg is kept intact, and a weight (blue square) is attached. **A**_**2**_. When a weight (blue arrow) is added, the additional gravitational force is sufficient to elicit a measureable change in leg posture upon inactivation of the motor neurons. **B.** The two camera views are used to reconstruct the 3D configuration of the leg. The cross placed on the fly’s back (blue cross) is used to align the leg with respect to the body. **C**. Multiple trials for each body orientation is shown. Body was rotated through five angles from -30 to +30. Each body orientation is shown in different color. Same color scheme will signify body orientation in Figures 1-3. For each orientation, the leg configuration is similar. The leg configuration changes with changes in body angle because the gravitational torque is different.

To increase gravitational force on the leg, we added a weight (∼20-100x the leg mass) to the tarsus (Figure 1A, Supplementary Video 1). Adding a weight resulted in much larger changes in limb posture upon inactivation. Two camera views were used to determine the three-dimensional configuration of the leg, and the tether placed on the fly’s back was used to determine the leg configuration in a body-centered coordinate system (Figure 1B). There was a change in posture with changes in the angle of body rotation (Figure 1C). The legs rest in distinct and consistent postures that are strongly affected by the angle of the body (Figure 1C) implying a relationship between limb posture and passive forces.

At each body angle, we inactivated the motor neurons multiple times, measured the joint angles and the corresponding gravitational torques to determine the relationship between joint angle and passive torques. We hypothesized that, for small deviations from the resting position, the passive torques can be modeled as an angular spring [3].. Figure 2A illustrates this idea for a 1-segment limb; the passive force is conceptualized as a spring which has a rest angle θ_0_ (red dotted line) at which there is no passive force. Gravitational forces acting on the weight will result in a deviation from θ_0_ which in turn would result in restorative forces that increase linearly with the angular deviation from rest angle. The joint will rest at an angle θ at which the torque due to restorative forces cancel those due to the gravitational forces. Gravitational torques can be calculated from the limb configuration, allowing us to infer the passive torque, which for a linear spring is given by the product of the angular spring constant and angular deviation from rest angle.

**Figure 2.**
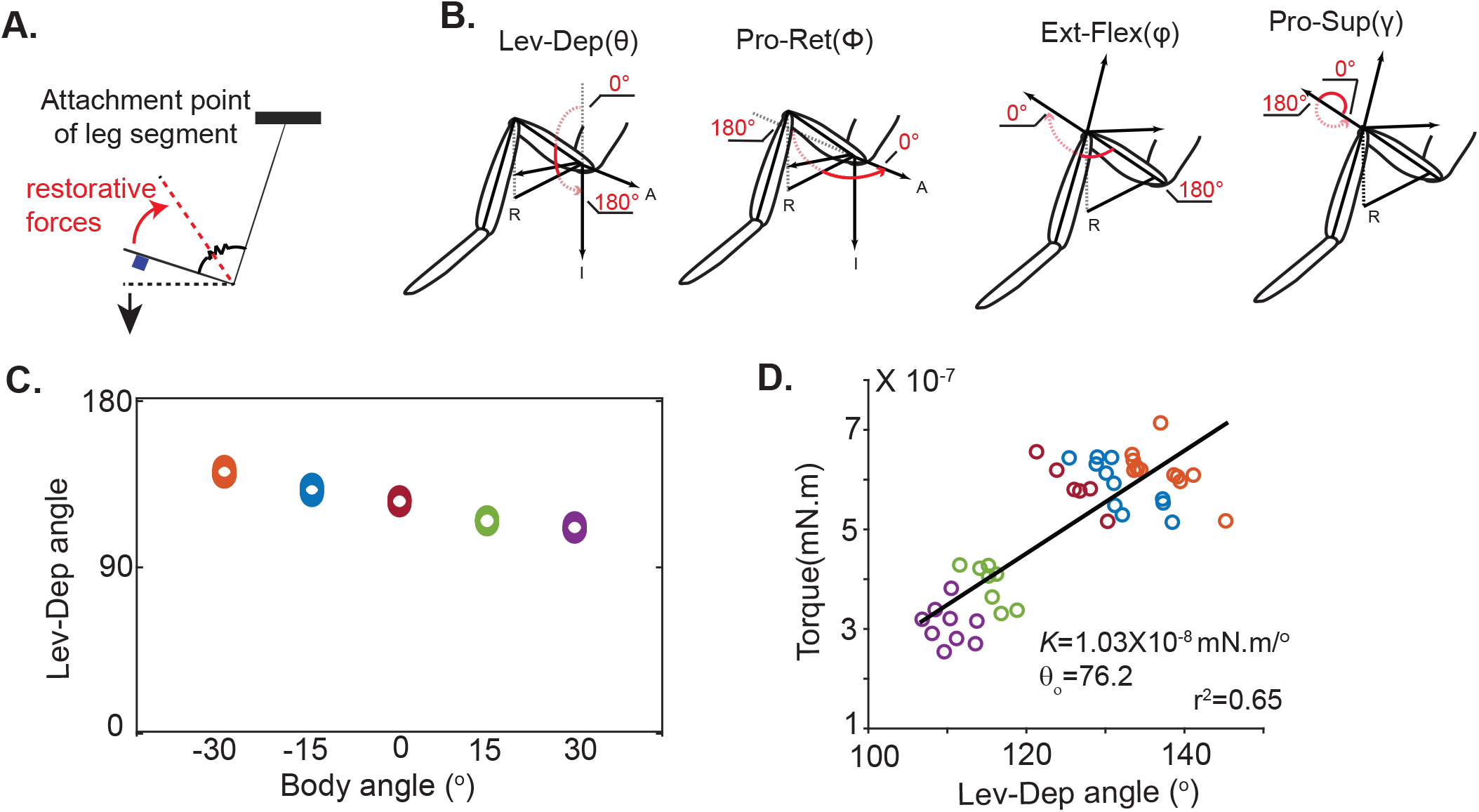
An angular spring describes the passive torques at each joint. **A**.Schematic that describes our analytical framework. As gravity (black arrow) pulls a limb segment away from its rest position (dotted red), passive forces increase linearly until torques due the passive forces balance torques due to gravity (at black dotted line). **B**. Definition of each of the four degrees of freedom (R=Right, A=Anterior, I=Inferior). **C**. Equilibrium angle is plotted as a function of body angle and changes with body rotation. Each marker is a separate measurement (n >6). As in Figure 1C, marker color denotes body orientation. **D**. The gravitational torque around the levation-depression axis was calculated. The torque changes linearly with the joint angle as shown by the large correlation between the levation-depression angle and the passive torque generated to counter gravitation forces.

Because the legs have multiple degrees of freedom, although conceptually like a single joint, the calculation of torques about each axis is somewhat more involved (see Methods). Calculation of the gravitational torque requires estimating the limb configuration and aligning leg configuration to the vertical. We consider 4-degrees of freedom: an insect thorax-coxa (*ThC*) joint has two-degrees of freedom (protraction-retraction and pronation-supination); the levation-depression degree of freedom is at the coxa-trochanter (*CTr*) joint (Figure 2B). There is an additional degree of freedom at the *FTi* joint. We estimate the torque about each degree of freedom.

The method for calculating passive torques is shown in Figure 2C and 2D using the example of one joint – levation-depression – in a single fly; group data is presented in Figure 3. As the body angle is changed, the leg stops at different levation-depression angles (Figure 2C) because the gravitational torque about the levation-depression joint changes with body orientation. If passive forces can be approximated as a linear spring, we anticipate a linear relationship between torque and angle. This linearity is exactly what we observe (Figure 2D). The spring constant *K*_θ_ and rest angle θ_0_ was calculated using linear regression (Figure 2D, black line). The spring constant *K*_θ_ showed some variability across flies as did the rest angle θ_0_ (Figure 2D).

**Figure 3.**
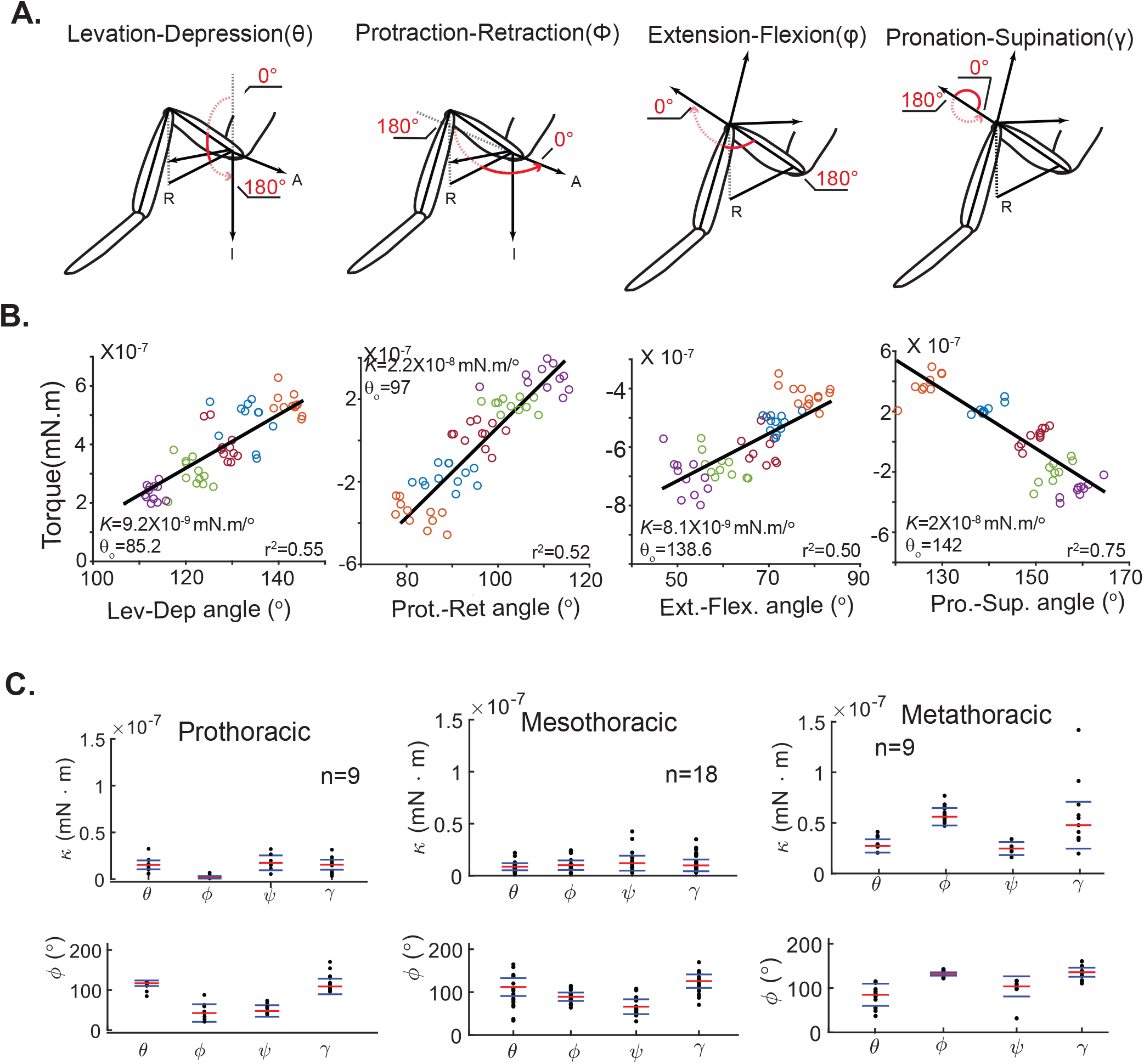
Passive torques at each of the four joints within a leg are well-described by a linear spring A. Definition of each of the four degrees of freedom (R=Right, A=Anterior, I=Inferior). Same as Figure 2A. **B**. Torque-angle relationships for the joints in the mesothoracic leg showing a linear relationship for each degree of freedom. Number of trials between 9-12. **C**. The passive force spring constant (κ) and equilibrium joint angles for four degrees of freedom for each leg. The red line is the median; the blue lines mark the inter-quartile range. The median values can also be found in Table 1.

The rest angle at a given rotation showed some variability; the standard deviation for the data shown in Figure 2D is ∼10°; this variation is consistent across most flies. To better understand the sources of this variability, we quantified the contribution of different sources of variability: We first considered the variability due to reconstruction errors. To this end, we evaluated errors due to calibration and used the calibration errors to estimate differences in both the joint angles and the sensitivity of the estimated *K* to these changes and found that calibration errors are small as the interquartile range is <5% different from the reported value (Figure 2-S1). Next, we focused on biological reasons for variability. Because muscles also receive octopaminergic input, it is possible that the variability is due to different levels of octopaminergic modulation in different trials. To address this issue, we performed the same experiment in flies where both the octopaminergic and glutamatergic neurons are silenced (Figure 2-S2). We found no difference in variability when both neural populations are inactivated (Figure 2-S2). Experiments using a larger mass resulted in a decrease in variability but did not yield different values for *K* (Figure 2-S2). Overall, a larger mass did result in lower variance, but it also caused the FTi joint to be extended to its limits in some cases. The decreased variability with higher weights does provide hints to the main sources of variability: Joint friction and coupling between different degrees of freedom mean that the gravitational torque acting on a joint will not bring it to rest at a unique position. A full analysis is beyond the scope of this study, but we have performed some analysis and discussed the implication (Figure 2-S3, methods and discussion).

We measured the passive forces at each of the other joints (Figure 3A). For each degree of freedom, we observed a linear relationship between torque and angle allowing us to calculate the angular spring constant and the rest angle (Figure 3B). The equilibrium angle and spring constant for each joint on each of the three right legs were calculated (Figure 3C and Table 1).

**Table 1:** Median stiffness for each of the measured joints in mN/^º^.

|  | Lev-Dep | Ret-Pro | Ext-Flex | Pro-sup |
| --- | --- | --- | --- | --- |
| Prothoracic | $1.5 \times 10^{-8}$ | $1.9 \times 10^{-9}$ | $1.7 \times 10^{-8}$ | $1.5 \times 10^{-8}$ |
| Mesothoracic | $8.6 \times 10^{-9}$ | $1.1 \times 10^{-8}$ | $2.5 \times 10^{-8}$ | $1 \times 10^{-8}$ |
| Metathoracic | $2.7 \times 10^{-8}$ | $5.6 \times 10^{-8}$ | $1.7 \times 10^{-8}$ | $4.7 \times 10^{-8}$ |

The interquartile range of spring constant implies about a 2-fold range across flies (Figure 3C). With some exceptions, joints within a leg have similar stiffness: The most significant exception is that the retraction-protraction stiffness is smaller in the prothoracic legs compared to other joints in the same leg, and larger in comparison to the other joints in the metathoracic leg. There is a corresponding difference in the joint angle with the prothoracic leg being more protracted and the metathoracic leg being more retracted compared to the mesothoracic leg.

There is more difference in spring constant across legs. The spring constant for the prothoracic legs are about 50% larger than the stiffness of the mesothoracic legs except for the retraction-protraction axis whose spring constant is much lower. The spring constants for the metathoracic leg is much higher.

How strong are the passive forces in flies? One method to contextualize the passive forces is to assess whether these passive forces are strong enough to support the fly. To assess how the measured passive forces compare with the forces necessary to support the fly’s weight, we created a model of the fly using the physics simulation engine OpenSim[18] . The fly was modeled based on the measured dimensions of the fly’s head, thorax, and abdomen, and each of the legs (Table 2). (Figure 4A and Supplementary video 2). The ThC joint was modeled as a ball joint with three degrees of freedom, and the FTi joint as a hinge joint with one degree of freedom. We incorporate the CTr joint movement as the third DOF in the ThC joint. We did not measure passive torques at the TiTa joint, and thus the CTr and TiTa joints are both kept fixed at their equilibrium positions in the model (Figure 4A and Supplementary video 2).

**Table 2.**
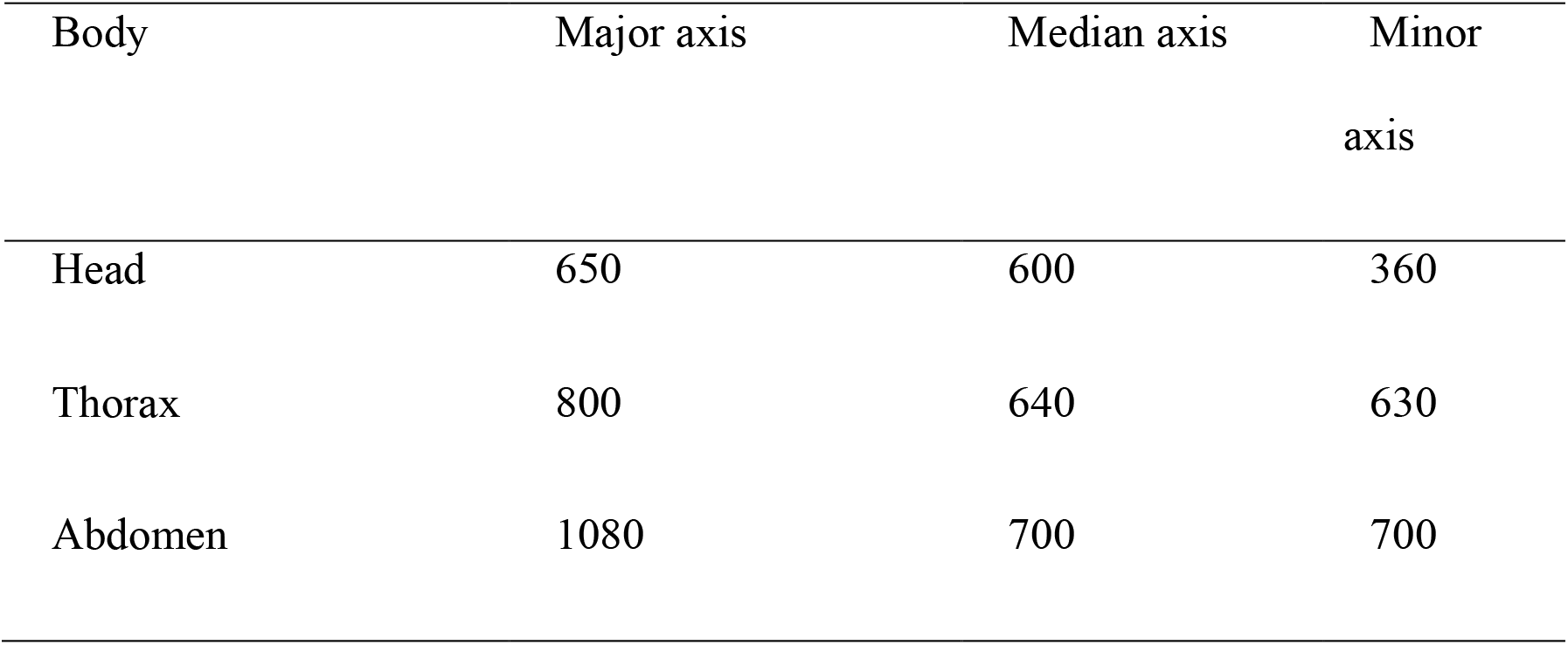
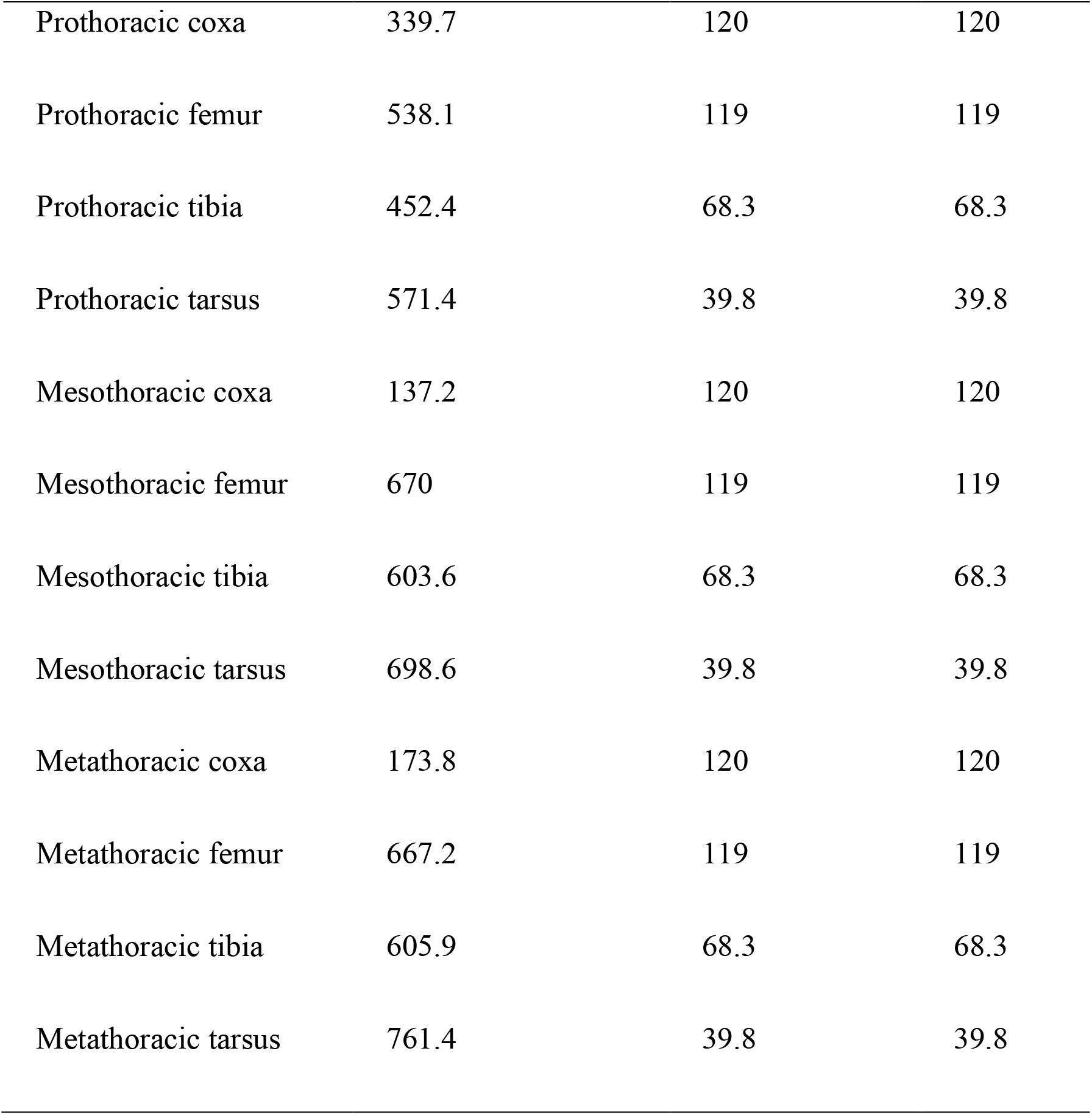
Ellipsoid radii for each body (μm)

| Body | Major axis | Median axis | Minor axis |
| --- | --- | --- | --- |
| Head | 650 | 600 | 360 |
| Thorax | 800 | 640 | 630 |
| Abdomen | 1080 | 700 | 700 |
| Prothoracic coxa | 339.7 | 120 | 120 |
| Prothoracic femur | 538.1 | 119 | 119 |
| Prothoracic tibia | 452.4 | 68.3 | 68.3 |
| Prothoracic tarsus | 571.4 | 39.8 | 39.8 |
| Mesothoracic coxa | 137.2 | 120 | 120 |
| Mesothoracic femur | 670 | 119 | 119 |
| Mesothoracic tibia | 603.6 | 68.3 | 68.3 |
| Mesothoracic tarsus | 698.6 | 39.8 | 39.8 |
| Metathoracic coxa | 173.8 | 120 | 120 |
| Metathoracic femur | 667.2 | 119 | 119 |
| Metathoracic tibia | 605.9 | 68.3 | 68.3 |
| Metathoracic tarsus | 761.4 | 39.8 | 39.8 |

**Figure 4.**
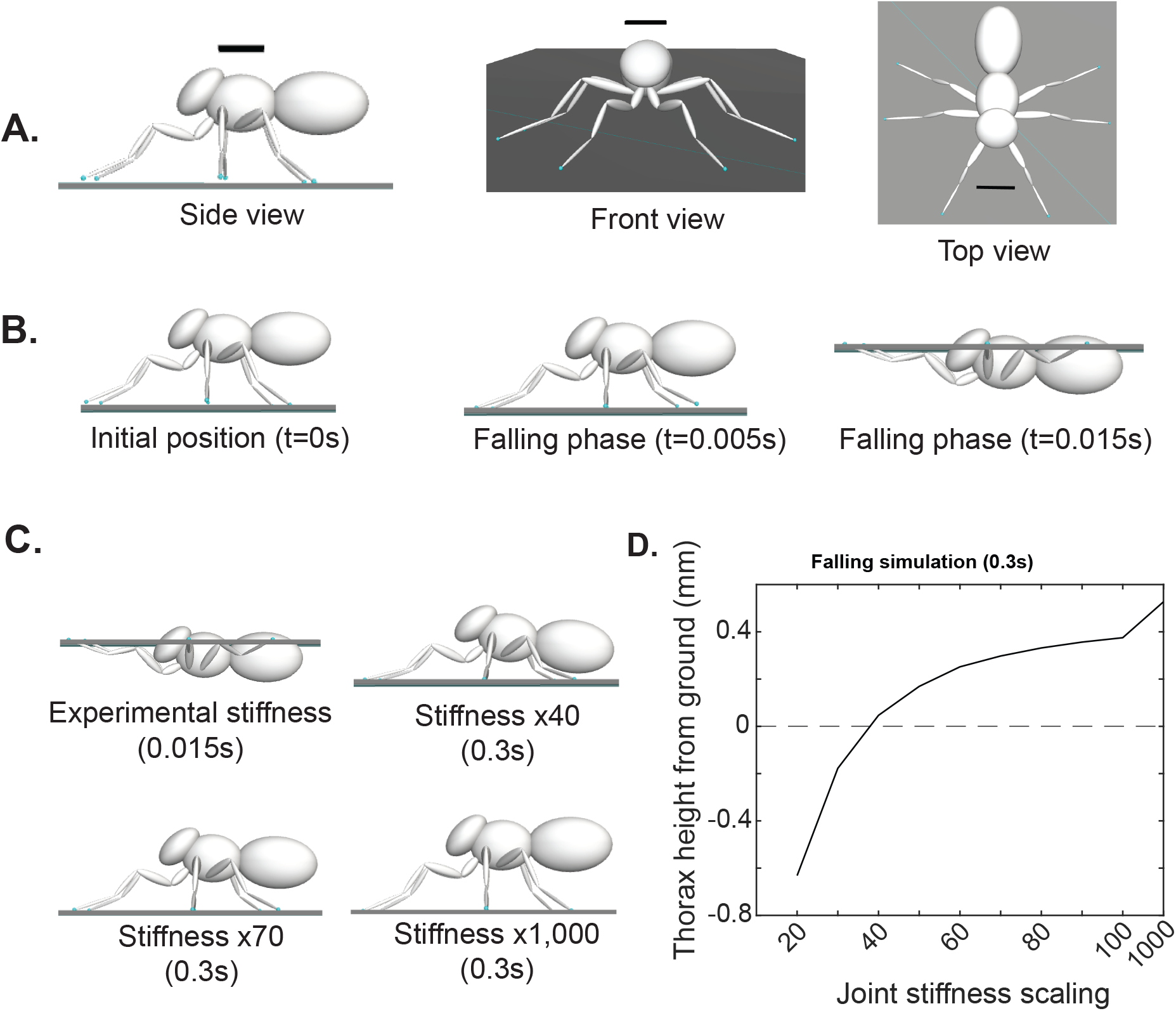
OpenSim model shows that the passive forces are ∼100-fold weaker than the forces required to support the fly. A. Schematic of the fly in the OpenSim model showing the fly in all three views (Scale bar = 0.5mm). **B**. Three timepoints of the simulation showing that the fly falls within 20 millisecond. **C**. Final equilibrium positions of the body at 4 different stiffnesses. **D**. Height of the CoM as a function of joint stiffness. The x-axis is scaled so that measured stiffness is 1.

Flies simulated with the measured passive stiffness were unable to support themselves and fell within 20 milliseconds, which is close to the freefall time (Figure 4B), implying that the weight of the fly is much larger than can be supported passively. As we increased the passive stiffness, we found that a 40-fold increase implemented uniformly across all leg joints was necessary to support the fly. At that force, the fly was barely supported over the ground (Figure 4C). As we increased the stiffness further, a more normal posture like the experimental flies was observed. Based on the height of the fly above the ground, we estimate that the stiffness is ∼80-fold higher than the measured passive forces. The plot of equilibrium height and joint stiffness is shown in Figure 4D.

To test the prediction made by the OpenSim model, we placed the fly of the same genotype as the ones used for tethered experiments - *VGlut-Gal4; UAS-GtACR1* - in a small rectangular chamber and video-recorded multiple views of freely walking flies as we inactivated the motor neurons with a green light (Figure 5A). We selected inactivation trials in which the flies are standing on all six legs, i.e., they are stopped. Using DeepLabCut[19] (DLC), we automatically tracked a point on the fly’s head in all views. Using these views, we performed a transformation to reconstruct the 3-dimensional position of the fly (see methods for details). We found that inactivation of motor neurons resulted in a rapid decrease in the fly’s height. The fly fell to the ground on every trial (n=92 from 8 flies). One trial is shown in Figure 5B; in this trial, the fall initiates in ∼40 milliseconds, there is a rapid drop – during which some part of the fly’s thorax or abdomen touches the ground. In some trials, the rapid drop is followed by a second phase – during which the height of the head continues to change as more of the body collapses to the ground.

**Figure 5.**
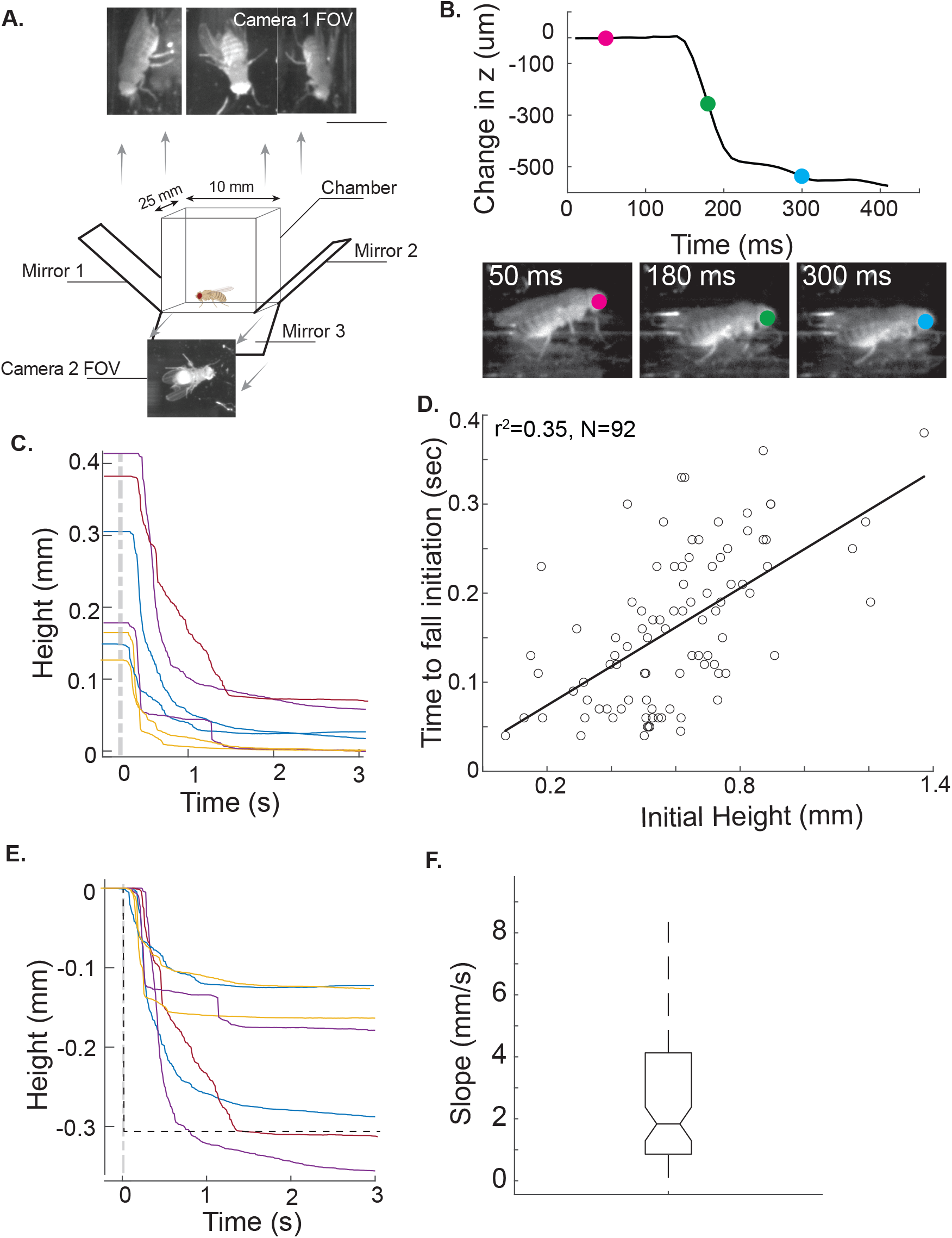
Passive forces alone are not stong enough to support the weight of a fly. A. A schematic of the walking chamber used to collect data about falling experiments. Because we have four views of the fly, the fly’s head could be located in multiple views to extract its position in all three dimensions. **B**. An example of a falling trial. Light on at t=0. Three frames showing the position of the fly at different times is shown. **C**. Flies stand at different heights in different trials. Few trials from one fly are plotted - the starting height varies from 0.4 mm to 0.1 mm in this fly. **D**. Time it takes to initiate a fall depends on the height of the fly. There is a significant correlation between height and time for fall initiation (r^2^=0.35). **E**. Apart from the time to initiate a fall, the rate of falling is much slower than that anticipated from immediate dissipation of the active forces (dotted line). **F**. The distribution of the slopes of the fall. Note that the rate of falling if the active forces dissipated immediately would be 37 mm/s.

As expected from the measurement of passive forces, the passive forces are too small to support the fly’s weight. However, there were two unexpected findings. The first finding is that there was a large variation in the initial height of the fly (Figure 5C), consistent with a recent study of flies walking on a treadmill[20]. This large variation in initial height implies that the active forces supporting the flies vary from trial-to-trial both within the same fly and across flies. Based on the OpenSim model, an active force of only 40X the passive force is necessary to support a fly standing at a height of 0.1 mm; the required force increases to 100X for a height of 0.4 mm (see Figure 4D). Consistent with this higher active force as the fly’s height increases, the time it takes to initiate a fall also increases with the height of the fly (Figure 5D). The time to initiate a fall varies from 40 milliseconds to 300 milliseconds, partly depending on the height of the fly.

Second, although the flies fell on each trial, it took much longer for the actual flies to fall compared to the simulated flies. As discussed above, one component of the difference is the time it takes to initiate a fall. A second component is the rate at which the fly fell to the ground. The rate at which the fly fell to the ground was much smaller in the experimental flies than it was in the simulated flies (Figure 5E). The median rate of falling was 1.3 mm/s compared to 37 mm/s for the simulated flies (Figure 5F).

The most likely reason for the longer than expected time for the fly to fall are delays associated with motor neuron inactivation and muscle inactivation. Based on measurements using the tethered fly (such as the one in Supplementary Video 1), we estimated that it takes about 350 milliseconds for the active forces to decay based on the time it takes for a fly’s limb to reach the new limb configuration defined by passive forces alone. To further estimate the possible delays, we performed in vivo whole-cell patch-clamp recordings from motor neurons. We used a line (e49-Gal4) in which it is easier to identify the motor neurons because of their position in the VNC as opposed to the broader set of neurons labeled by VGlut-Gal4. Using a digital mirror device (DMD) that allows us to stimulate either the entire VNC or only the recorded motor neuron, we show that the inhibition of motor neuron is intrinsic, i.e., reflects opening of ion-channels in the motor neuron itself (Figure 6B). A spot of light on the cell body produces as much of the hyperpolarization as stimulating the entire fly (mean of 11.3 mV vs 13.1 mV across 9 neurons). Conversely, excluding the cell body produces only a small effect on the MN (mean of 2.6 mV). The normalized change in voltage as a function of time is shown in Figure 6C. The MNs had a similar time-course: the initiation of inhibitory response took ∼30 milliseconds; this number is similar in all MNs (Figure 6C). The time it took for MNs to complete 90% inactivation ranged between 52-114 milliseconds (Figure 6C) implying that passive forces will take at least between 50-100 millisecond to decline as this measurement does not consider the delay between motor neuron inactivation and loss of muscle stiffness. This delay – between motor neuron inactivation and loss of muscle stiffness - will be different for different muscles and estimates for many muscles are not available currently. The delay for the slow tibial motor neuron is ∼40 millisecond[21]. Assuming a similar delay for all muscles, we reasoned that the decay of active forces will occur with a time constant equal to the sum of the two time constants discussed above giving a range of 100-150 milliseconds.

**Figure 6.**
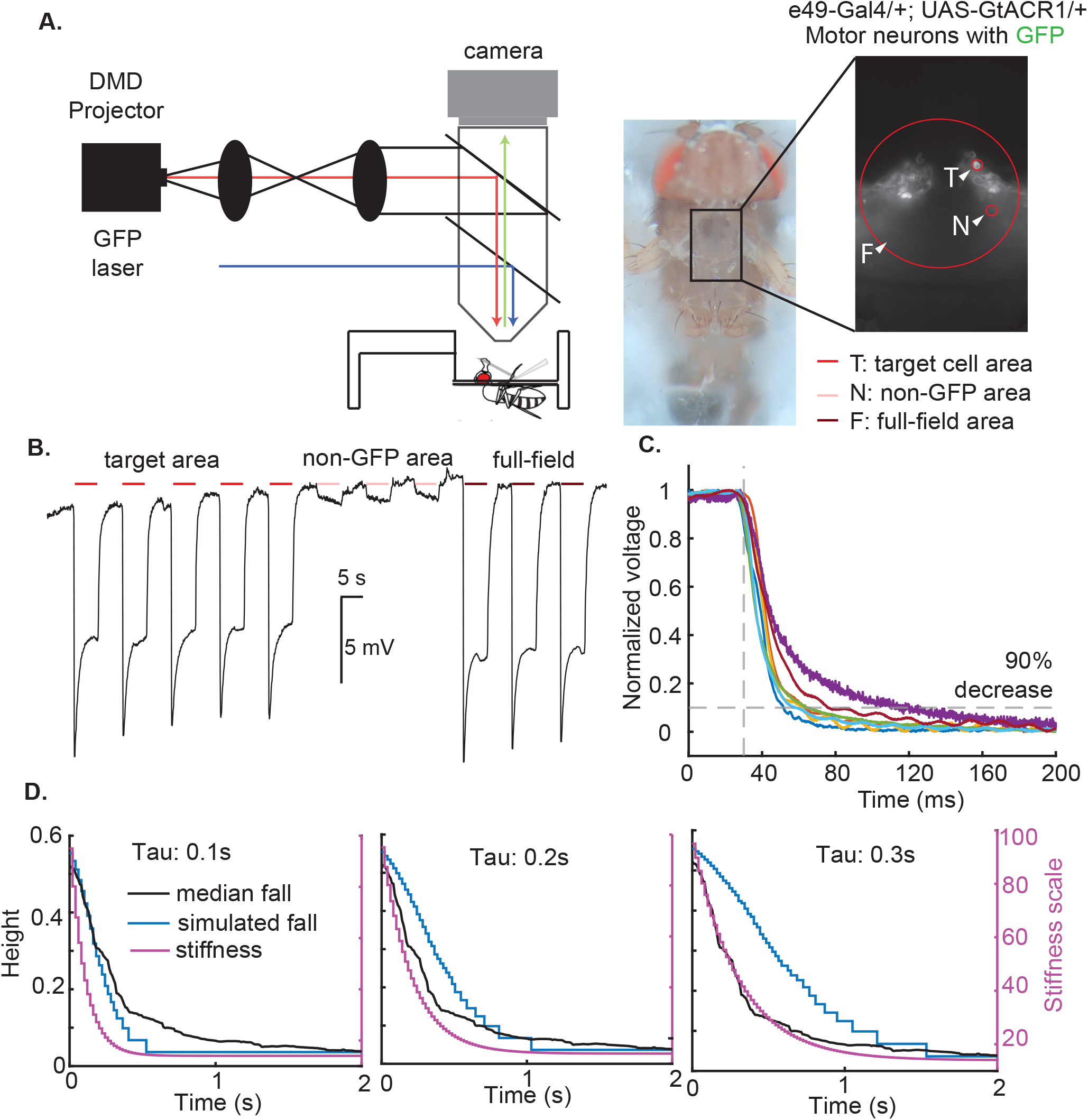
The dynamics of the fly’s fall suggest that active forces decay with a timeconstant of ∼100 ms and is consistent with the time-course of motor neuron inactivation. A. Schematic of the set-up used to evaluate motor neuron inactivation. A DMD projector was used to either shine light on the motor neuron cell body (T), stimulate a non-target area (N), or provide full-field stimulation (F). **B**. Sample trace from a single-neuron showing that stimulating just the cell-body (left 5 trials) causes similar inhibition as the full field stimulation (right 5 trials). Stimulating regions not including the cell body (middle 5 traces) causes only a small depolarization. **C**. Time course of hyperpolarization in different motor neurons. **D**. OpenSim simulations show that an exponential decay with a time constant 100 ms best fits the dynamics of the fall. An exponential decay of 200 ms and 300 ms produces falls that are considerably slower.

To estimate whether the temporal delays associated with neural inactivation and muscle inactivation can explain the time course of the fall that we observe, we created an OpenSim model in which the stiffness decayed exponentially. For the median fall, the fly stands at a height of 0.5 mm. Using the relationship plotted in Figure 4D, the active forces necessary to support the fly at that height is ∼ 90X the estimated passive forces. If we decrease the active forces such that the force at any time t is given by an exponential decay *f*(*t*) = *f*_*a*_ · *e* ^−*t*/τ^ + *f*_*p*_. Here *f*_*a*_ and *f*_*p*_ are active and passive forces, respectively. The τ represents the time constant of the decay of the forces. When the active forces decay at an exponential rate of 100 ms, the initial slope of the resulting simulated decay is similar to the median fall (Figure 6D); a rate of decay of 200 ms produces a fall that is considerably slower than the actual fall (6D). This simulation shows that dynamics of the fall are consistent with the expected rate at which the active force would decay.

## Discussion

By leveraging the genetic tools in *Drosophila*, we could measure passive torques in all the joints of the fly’s middle leg except the tibia-tarsus joint within an intact leg. This method complements more invasive methods used in larger insects [22] by providing an estimate of passive torques for all leg joints in a completely intact leg. Optogenetics also allows for reversible inactivation, and measurements in a completely intact animal which is not possible with denervation techniques employed in large insects. A possible limitation of this method is that it is not as informative regarding the origin of the passive forces; a limitation that can be overcome in future work. As demonstrated in this study, the usefulness of this method is in connecting the measured passive torques to its consequences for the whole animal.

There are several important differences between our passive measurements, and those made previously. In most previous studies, a single joint is studied in isolation, and therefore there is a single degree of freedom. In this study, we examine the entire leg with multiple degrees of freedom. When the motor neurons are silenced, and the leg posture changes as it finds its new equilibrium, this new position can be different on different trials because of the multiple degrees of freedom. For instance, the torque along a given axis of rotation need not only be controlled by the angle of rotation about that axis, but also by other angular degrees of freedom relevant at the given joint. Therefore, one can achieve the same torque with slightly different angle of rotation about the torque axis due to compensatory changes in other angular degrees of freedom. This implies that equilibrium can be reached not just at a unique angle of rotation but rather for a small range of angles.

Despite this degeneracy, the passive torques in each joint is well described as a linear spring over a relatively large range of motion that extends over 40 degrees on each side of the rest position (see Figure 3). This linearity is consistent with the linearity in passive forces estimated during aimed leg movements in the locust [3]. Similarly, measurements in the FTi joint of a stick insect [6] show a roughly linear dependence on angle except when the joint is extended beyond 150°. The linearity observed here might reflect a similar cancelation between two non-linear processes as observed in the locust and stick insect: The non-linear increase in the force exerted as the joint moves away from its rest angle which in counteracted by a similar non-linear decrease in the moment arm.

The median stiffness of all the joints we investigated lie within a 7-fold range from 8X10^-9^ Nm/° for the levation-depression in the mesothoracic leg to 6X10^-8^ Nm/° for the retraction-protraction in the metathoracic leg. Most joints within a leg have similar stiffness; larger differences are observed across legs: prothoracic legs are stiffer than mesothoracic legs and methathoracic legs have the stiffest joints. Although more work is needed to fully understand differences across joints and legs, the narrow range of stiffness values are consistent with the similarity in the number of muscle fibers at each joint [23]. Differences in moment arm due to differences in attachment point are more likely to underlie differences across stiffness values than differences in the number of muscle fibers [24]. The most consistent difference in the rest angle is the difference in the retraction-protraction angle between the legs with the prothoracic leg more protracted and metathoracic leg more retracted than the mesothoracic leg. Similar differences were observed in the passive position of stick insect legs, and these differences were hypothesized to occur due to the differences in muscle attachment and resulting moment arms [24].

The range of stiffness values for a given joint across flies has an interquartile range that suggests about 30% variation on either side of the median. The contributions to this variability from errors resulting from the errors in tracking, in 3D-reconstruction, and uncertainty in linear regression is small (Figure 2-S1). Therefore, much of this variability is biological and comparable to measurements made in larger animals such as locust and stick insect [5, 6]. We also did not find a strong correlation between joint stiffness at the different joints in the same fly; higher stiffness at one joint is not predictive of higher stiffness at another joint further suggesting that small differences in spring constants are probably not critical.

In many studies, an asymmetry is observed between the passive forces when the joint angle is increased versus when the angle is decreased resulting in about a 10° difference in rest position in locust [4], stick insect [1], and false stick insect [5] depending on which direction the approach to rest is made. This asymmetry results from the different cross-sections of the muscles as well as different origins of the passive forces in the muscle versus at the joint. The data in this study is more consistent with symmetry between extension and flexion because the spring constants appear to be similar irrespective of whether the joint is more extended or flexed. One reason for the symmetry might be the number of muscle fibers are similar between extensors and flexor.

Another reason is that in larger insects, there are a group of inhibitory motor neurons that play a crucial role in tuning passive muscle properties, and these properties could be different whether the muscle is returning to the rest position from an extended or a flexed position [25]. In flies, these inhibitory motor neurons are not present [21].

Previous estimates of passive forces in the FTi joint have suggested that they might be sufficient to support the weight of the insect. As an example, the FTi joint on each leg of a stick insect produces 6 mN of passive force [1]; together the 6 legs of the stick insect should produce forces sufficient to support the weight of a stick insect (10 mN). Similar estimates in cockroaches suggested that a mere 20° change in the FTi joint is enough to generate sufficient passive forces to support a cockroach. Based on these findings, and the fact that the scaling laws should make passive forces even larger in relation to the weight of the insect, we expected that the passive forces should be able to support the weight of a fly. Surprisingly, the passive torques we measured were insufficient to support the weight of the fly. In fact, the required torques were approximately two orders of magnitude too weak. Both the predictions from our measurement with tethered flies, and experiments with freely walking flies support these conclusions. Back of the envelope calculations based on previous measurements suggest that stick insects, locusts, cockroaches should all be able to support their weight using passive forces alone; at the very least, these forces are unlikely to be 100-fold smaller. Thus, it appears that the passive torques measured in flies are at least two-orders of magnitude smaller than anticipated from measurements in larger insects. This difference is particularly striking because we are likely measuring the dynamic passive forces that are present soon after movement that are much larger [26-29], rather than the static forces measured typically.

More work is necessary to assess whether this discrepancy is due to methodological differences or reflect real biological differences. There are key differences between the genetic inactivation method employed in this study, and denervated or isolated legs employed in studies in larger insects. In the genetically inactivated flies, other sources of modulation remain preserved that might contribute to the measurement. Denervated/isolated leg preparations provide measurements that are closer to the actual mechanical properties of the leg. One possibility is that passive torques scale with the moment of inertia of the leg. If that is the case, the corresponding scaling law would be (mass)^5/3^ implying that the passive torques would scale non-linearly with the mass of the insect. This would neatly explain why passive forces and the resulting torques produced in locust and stick insect which are 1000-fold heavier than flies are in the correct order of magnitude to support the weight of these insects while being 100-fold weaker in flies since the ratio of passive forces to mass scales by (mass)^2/3^.

Even though passive forces are not strong enough to support the fly, they are strong enough to impact both the posture of the limb and its movement. Femur has a mass of ∼7 μg, and length of 600 μm yielding a moment of inertia of 2.6X10^-15^ Nm. The passive torques produced by deflecting the femur by 10° is ∼10^-10^. Thus, the passive torques produced are several orders of magnitude greater than the gravitational torques on the femur, and account for the gravity independent positions of the limb. Passive forces will also have a large effect on the movement of the limb.

A challenge with motor control in insect limb is that the forces required are small when compared to forces produced by muscles and joints within these limbs. To achieve these small forces, active forces much larger than the forces necessary to move the limb are counteracted with equally large passive forces leaving a small residual force[3]. This finding has led to the conclusion that passive forces in both locust [3] and stick insects [1] are of the same magnitude as the active forces. There is some evidence to suggest that passive forces are specifically adapted to match the active forces [5]. However, passive and active forces have only been compared during movements when the leg is unimpeded by ground contact. Our results suggest that the active forces are ∼100-fold larger than the passive forces during weight bearing tasks. It is possible that much smaller active forces are generated once the leg is lifted off the ground, and that the active forces during unimpeded ground contact in *Drosophila*, too, is of the same order as the passive forces. Alternatively, it is possible that flies might operate in a different force regime in which the passive forces are much greater than the residual torque, and the active force itself is much greater than the passive forces.

Another issue to consider in the motor control of fly limbs is that the decay of active forces will take time. Cessation of firing, decrease in forces following decrease in firing would require ∼100 millisecond which is of the same order of magnitude as a single step in the fly which appears to be implausible. It is more likely that fly locomotion works in a regime in which active forces cancel each other out to produce small residual forces.

## Methods

### 1. Flies

All experiments, except motor neuron recordings, were performed on 3-5 day old female flies of genotype *VGlut1(OK371)-Gal4; UAS-GtACR1*. Motor neuron recordings were performed on *e49-Gal4; UAS-GtACR1* flies. The flies were reared at 25°C, and 12 hr:12 hr light: dark cycle until eclosion when they were transferred to food containing retinal (0.02% concentration of all-trans retinal) and kept in complete darkness until experiments were performed. We also performed experiments silencing both octopaminergic and glutamatergic neurons (Figure 2-S2) using *VGlut1(OK371)-Gal4/Tdc2-Gal4; UAS-GtACR1* flies.

### 2. Experiments on tethered flies

Flies were attached to a tether using a small drop of UV glue (Kemxert-300) that was cured using UV light provided by a UV-gun (LED-100 from Electro-lite). We made the tether using a D-Sub pin (Digi-Key #A2160-ND) with a 1cm length hypodermic tube (0.025”OD x 0.017”ID) epoxied on the female end. Two pieces of tungsten wire (0.005”OD), 1cm x 0.5cm epoxied together at a 90-degree angle made up the long and short axis for the tether, and this was attached inside the exposed end of hypodermic tube with epoxy. The tether was mounted to a precision rotational mount (Edmund Optics #66514) to allow small, precise rotations of the fly. In pilot experiments, we found that sometimes the fly’s legs would get entangled with each other when they are inactivated preventing an accurate assessment of the limb position. Therefore, in most experiments all legs except the leg being experimented with was removed. For tethered experiments, all data was collected with a weight attached to the leg. Most experiments were performed with a weight whose mass was ∼20X the mass of the fly’s rear leg (a 1/4mm length piece of wire was weighed to be 0.19 mg), then attached to the distal tarsus using a small drop of UV glue cured with a UV light). We also performed some experiments with weight that was ∼50 X and 100X the mass of the rear leg.

The flies were imaged using two USB 3.0 camera Basler acA1920-150 μm (50hz, 1920 px x 1200 px) and a Infinistix lens (0.5X, Edmund Optics #54675) to collect videos from two vantage points. A filter was placed on the lens to block visible light (570nm long pass filter, Edmund Optics #54656). Exposure was set at 1 millisecond. The flies were illuminated with 850 nm high-powered LED (Thor Labs #M850LP1). Each trial started with a 60-second period during which the neurons were not inactivated. During the rest of the trial 8 seconds of inactivation were interleaved with 2 second periods of rest. During the stimulus period, the flies were irradiated with green light (Thor Labs #M530L3).

Passive stop frames were automatically extracted from the two videos by assessing frames during which the legs were completely stopped during the inactivation period. This extraction method is not foolproof as the legs can get entangled with the stubs of the other legs, and to the cross that marked the long and short axis of the fly . Therefore, we manually inspected all stop frames and eliminated these cases. All the data for mesothoracic leg was collected from 20X weights, the data from metathoracic leg was a mixture of 20X and 100X weights, and from the prothoracic leg was a mixture of 20X and 50X weights.

We annotated four points on the fly’s leg on each camera view: the coxa, FTi joint, TiTa joint, the location of the weight, and the end of the tarsi; using direct linear transformation[30], we used the two views to obtain the 3-D position of each of the four points. Because rotations are not pure rotation, and there is some chance of translation, we used a cross attached to the dorsal surface of the fly’s thorax as an internal reference frame; the two axes of this cross were aligned to the fly’s long and short axes.

Calculation of torques and joint angles is described in the next section. After these calculations, we performed a robust linear fit using MATLAB.

The tethered experiments were performed in three batches. In the initial experiments, we targeted 18-24 trials per orientation. In the second and third set we used 9-12 trials, since we realized that the data was less variable than we expected. Not every trial could be used. Trials were rejected because the legs either got entangled with other legs or with the cross. Some experiments had to be stopped before all the trials could be completed because 1) The weight fell off, 2) flies could sometimes ripoff all or part of the leg while attempting to remove the weight, 3) between each inactivation bout, the leg moved vigorously. Experiments were stopped if this movement ceased before the end of the experiment, 4) Finally, sometimes the fly’s leg could get permanently stuck. Regardless, we only used flies in which we had at least 6 trials for a given orientation.

### 2. Calculation of passive joint forces

This section describes the calculation of passive joint torques for each degree of freedom. We use the tracked world coordinates of physical wires oriented along the fly’s body axes to establish an anatomical coordinate system at the tether, which we will refer to as the body axes. The position vector for the left-right axis is defined by subtracting the right (*w*_*R*_) from the left (*w*_*L*_) world coordinate of the physical wire:

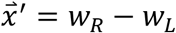

The position vector for the anterior-posterior axis is defined by subtracting the posterior (*w*_*P*_) from the anterior (*w*_*A*_) world coordinate of the oriented physical wire:

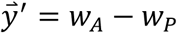

We then perform the Gram-Schmidt process for orthonormalizing a set of vectors to obtain orthogonal 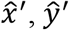 and 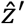 basis vectors. The left-right axis 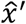 is defined by normalizing 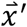:

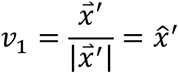

To obtain the orthogonal anterior-posterior axis, we first obtain the orthogonal projection of normalized 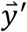 on the subspace spanned by 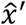:

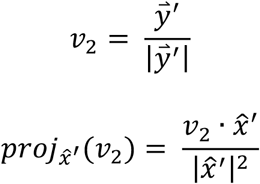

The difference between *v*_2_ and its orthogonal projection onto the previous subspace is orthogonal to the vectors in that subspace, thus giving us the 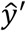 basis vector:

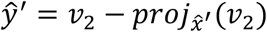

Similarly, to obtain the orthogonal dorsal-ventral axis, we first obtain the orthogonal projection of normalized 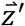 on the subspace spanned by 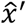 and 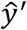:

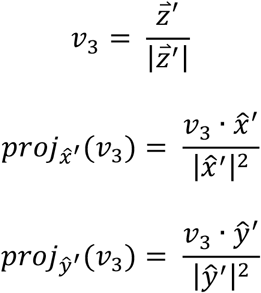

Likewise, the difference between *v*_3_ and its orthogonal projection onto the previous subspace is orthogonal to the vectors in that subspace, which yields the 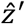 basis vector:

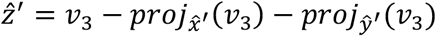

Each leg segment unit vector is defined as the radius vector relative to its point of attachment and are symbolized by 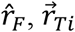, and 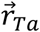 for the femur, tibia, and tarsus axes respectively. The femur vector is defined as the difference between the femur-tibia and coxa-trochanter joint positions in world coordinates:

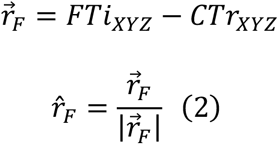

The tibia vector is defined as the difference between the tibia-tarsus and femur-tibia joint positions in world coordinates:

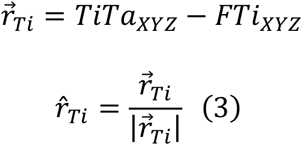

We use these unit vectors to calculate the four angles which describe the degrees of freedom considered here: Levation-depression (*θ*), protraction-retraction (*φ*), and pronation-supination (*γ*) at the proximal coxa joints, as well as extension-flexion (*Ψ*) at the femur-tibia joint.

The levation-depression angle (*θ*) is calculated as the angle formed by the femur’s projection in the body’s transverse 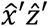-plane with the body’s dorsal-ventral 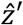-axis. In this convention, 0° corresponds to the femur pointing vertically. We obtain the femur vector’s components in the transverse plane:

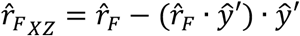

Finally, *θ* is the inverse cosine of the dot product between the femur’s transverse plane projection and 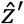:

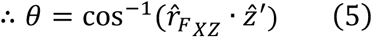

The protraction-retraction angle (*φ*) is calculated as the angle formed by the femur’s projection in the body’s coronal 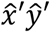-plane and the body’s anterior-posterior 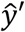-axis, with 0° corresponding to the femur pointing anteriorly.

We obtain the femur vector’s components in the coronal plane:

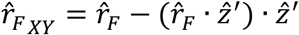

Finally, *φ* is the inverse cosine of the dot product between the femur’s *XY*-projection and 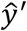:

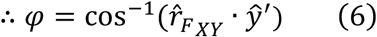

To determine angles at the femur-tibia joint we establish a reference frame at the femur vector 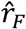. The femur-tibia (*FTi*) reference frame is defined by the orthonormal basis vectors 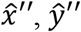, and 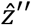 (Equations 7-9):

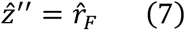

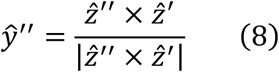

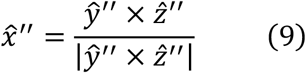

Using the femur-tibia reference frame, the extension-flexion angle (*Ψ*) is defined as the angle formed by the femur and tibia vectors (Equation 10):

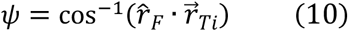

The pronation-supination angle (*γ*) is calculated by relating the elevation (*Ψ*) and azimuthal angle (*γ*) of 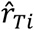 to the projection of 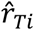 in the femur-tibia axes. We then solve for *γ* (Equation11):

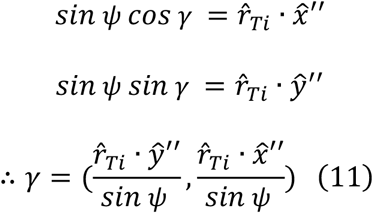

We then determine passive forces by evaluating torques along each rotational axis. The fly’s leg is static so it can be assumed that only gravitational and passive forces are acting on the leg. The gravitational torques due to limb mass are negligible with respect to the torque due to the applied weight at the tarsi (>20 times total leg mass), and thus they can be ignored. The gravity vector 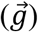 is defined with an acceleration of -9.81m/s^2^ along the vertical axis of the world coordinate system (*z*_*global*_). Equation 12 describes the net torque at the leg’s point of attachment, or origin, due to the applied weight 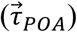 relative to the leg’s point of attachment. For the meso- and metathoracic legs, the point of attachment is defined as the coxa-trochanter joint position since the coxa is small and has a negligible influence on leg torque. For the prothoracic leg, it is defined as the thorax-coxa joint due to the much larger length of the prothoracic coxa. With *m*_*W*_ the mass of the applied weight, the net torque is thus represented by the cross products of the weight position vector and its gravitational force vector (Equation 12):

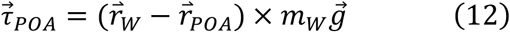

The torque vector at the femur-tibia joint 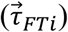 is calculated in the same way, but using the radius vector of the applied weight’s center of mass relative to the *FTi* joint (Equation 13):

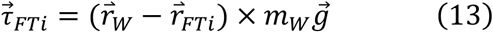

The levation-depression axis is represented by the vector orthogonal to the femur’s projection in the body’s transverse 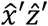-plane and the body’s dorsal-ventral 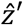-axis. Thus the torque τ_*θ*_ about this axis is the projection of 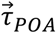 onto the unit vector formed by the cross product of 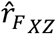 and 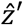 (Equation 14):

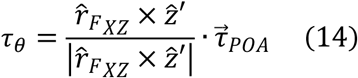

The protraction-retraction axis is represented by the vector orthogonal to the femur’s projection in the body’s coronal 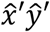-plane and the body’s right-left 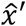-axis. Thus, the protraction-retraction torque (τ_*φ*_) about this axis is the projection of 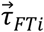 onto the vector formed by the cross product of 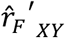 and 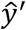 (Equation 15):

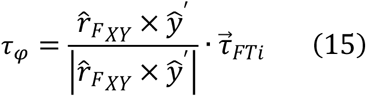

The extension-flexion axis is orthogonal to the femur 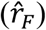 and tibia axis 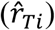. Thus, the extension-flexion torque (τ_*Ψ*_) is the projection of 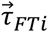 onto the cross product of 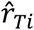 and 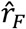:

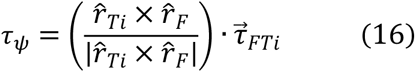

The pronation-supination axis exists along the femur vector represented by 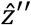. Thus, the pronation-supination torque is simply the projection of 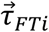 onto this vector (Equation 17):

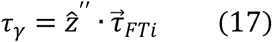

To verify the accuracy of our calculated joint parameters, we used forward kinematics and determined the mapping function from joint space to the world coordinate system in Cartesian space. Leg positions from each body rotation were reconstructed into world coordinates from joint positions and then overlaid onto the original tracking data. This verified that our angle calculations are capturing the passive stop positions accurately.

### 3. Estimating errors in the measurement of spring constant due to reconstruction

The accuracy of our 3D reconstructions is dependent on the quality of the Direct Linear Transformation (DLT) coefficients obtained from the two-camera system calibration. To verify that our DLT calibrations capture the transformation from 2D camera coordinates to 3D world coordinates, we used inverse DLT to obtain pixel coordinates from the 3D-reconstructed leg joint positions using corresponding calibration coefficients. We then overlaid these points on the corresponding frames for all acquisitions and found that the coefficients accurately capture the inverse transformation from 3D-reconstructed leg coordinates to corresponding pixel positions for each camera (Figure 2S1-A).

To estimate the error in the reconstruction attributable to the camera calibration, we performed the following procedure: First, we obtained a random subsample of half of the control points used in the calibration. The control points consist of the pixel coordinates of a fixed object for each camera, and its corresponding 3D coordinate. The control points were acquired by moving a motorized micromanipulator to desired locations. We then run the DLT calibration to obtain new coefficients, and then finally use this new calibration to predict the 3D coordinate of the other half of the control points. We repeat this procedure 100 times and record the error in the predictions for each iteration, which yields a distribution of errors along each axis for the camera calibration (Figure 2-S1B). We found that the error distribution for each camera calibration is well-described by a normal distribution, and that reconstruction error is low, on the order of <50μm for each axis (Figure 2-S1C).

We then performed error propagation to investigate the extent to which DLT error affects our findings. To do this, we performed 1000 simulations of error propagation for each 3D-reconstructed point by random sampling from the DLT camera calibration’s error distribution to obtain 1000 simulated legs. We used these simulated legs to measure the spring constant. We found that the resulting distributions of spring constants were in a small range (Figure 2-S1D and 2-S1E).

### 4. Evaluating the limitation of the linear approximation

Our data clearly indicates that passive forces can be approximated by a linear spring. However, the leg does not settle in the same position over multiple trials. One reason for this variation is that there are many degrees of freedom within the leg; the many degrees of freedom mean that the gravitational torque can cause displacement along any degree of freedom. One consequence of these multiple degrees of freedom is that the torque produced along a given degree of freedom is dependent on how the body is rotated. Rotating the body along its long axis causes a large torque along the levation-depression joint, and a small torque along the extension-flexion axis. The converse is true for rotation along the fly’s short axis.

Another consequence is that the variation in torque within the same experimental condition, i.e., body rotation. Some of this variation is because there is some coupling between different degrees of freedom (DOFs). A full analysis of this coupling is beyond the scope of this study. We performed some analyses to show that some of the variation comes from coupling from other DOFs.

To test whether joint torque variability correlates with actions at other DOFs, we obtained the residuals from our angle-torque fits and correlated these with the corresponding angles denoting the other DOFs. We found that in the majority of cases the analyzed residuals exhibit a correlation with angles at other DOFs, suggesting that residual torques for one DOF can be affected by movement at other DOFs (Figure 2-S3A). To further investigate this coupling, we performed regression using multiple combinations of angles to predict levation-depression torque and determine whether a better fit is obtained for the fly. We found that multiple regression results in a closer fit by plotting R^2^ vs. the combination of angle DOF predictors (Figure 2-S3B). The regression model for levation-depression torque (τ_*θ*_) thus can be described with retraction-protraction (*φ*) using the following regression model, where the passive torque equals the torque due to gravity (τ_*G*_) at equilibrium with τ_0_ the y-intercept and τ_*I*_ the torque due to other internal forces in the joint:

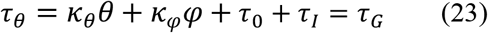

Using the stiffness coefficients κ_*θ*_ and κ_*φ*_ obtained from the multiple regression model, the effective angle due to *φ* thus becomes:

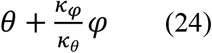

By plotting this effective angle vs. torque, we show that the fit is indeed improved due to the inclusion of the other DOF (Figure 2-S3C). Since the effective angles produced by the estimated coefficients from the multiple regression fits yield lower R^2^ values for LD torque than when only using the LD angle, it suggests that LD torque variation is partially explained by coupled actions at multiple DOFs.

### 5. Experiments in freely walking flies

Inactivation experiments to assess whether a freely-moving fly falls when its motor neurons are inactivated were performed in a walking chamber.

Flies were placed in a rectangular plastic chamber measuring 25mm × 10mm × 10mm (LxWxH) and left to acclimate for 10 minutes before starting the experiment. The flies were imaged using two USB 3.0 Basler acA1920-150 μm cameras (100Hz, 960 px x 520 px) equipped with telocentric lenses on both cameras (Edmund Optics 0.40x SilverTL, part number 56-677) to record the video at 100 fps at 960×520 resolution. A filter was placed on the lens to block visible light (720nm infrared filter, Hoya B-52RM72-GB). The flies were illuminated with 850nm high-powered LED (ThorLabs #M850L3). Trials were designed in the following way: 5 seconds of control conditions were followed by 15 seconds of inactivation and concluded by 5 seconds of control conditions. During the inactivation period, the flies were irradiated with green light (Thor Labs #M530L3). We collected 88 trials from 8 flies (with a minimum of 7 and a maximum of 12 trials per fly) where flies were located at the bottom of the chamber right before the onset of inactivation period.

The head of the fly was automatically tracked using DeepLabCut (DLC; Mathis et al., 2018). Briefly, the head was manually annotated on unique frames manually extracted from the recordings of the 8 flies within the dataset and we trained 4 distinct Resnet-50 (He et al., 2016) for the 2 mirror-reflected views, the central top view and bottom view.

To transform the two-dimensional camera coordinates from each of the four views into a single set of three-dimensional world coordinates, triangulation was performed. The position of a fluorescent microbead fixed at the tip of a pulled glass micropipette and moved using a micromanipulator was captured using a custom-made MATLAB interface (GUI). Briefly, images of the microbead at 130 known positions were taken and the position of the bead semi-automatically detected 3 times in the top camera view (in the 2 mirror reflections as well as the real chamber) and once in the bottom view. Triangulation was performed using direct linear transformation (DLT); the mean squared error was 49 μm.

The resulting transformation matrix was used to transform the 4 sets of head coordinates in the camera reference frame provided by the automatic DLC tracking to generate the fly’s head in world coordinates.

### 6. Simulation in OpenSim

To determine the degree to which our experimentally determined stiffness values at each degree of freedom are sufficient to support the legs, we incorporate these forces into a mechanical model of the fly and simulate this model dropping onto a platform with the force of gravity. The mechanical model derived from the fly’s anatomy (Figure 4A) was generated in OpenSim 4.1[18]. MATLAB integration modules were used to generate the model by defining rigid bodies interconnected by joints. The dimensions, masses, and center of mass positions for each body segment were obtained from previous experimental measurements and from earlier work[31]. Each body segment was approximated as an ellipsoid (dimensions in Table 2). To model the leg joint dynamics, we introduced a ball joint at the *ThC* with three degrees of freedom corresponding to protraction-retraction, pronation-supination, and levation-depression, as well as a hinge joint at the *FTi* with one degree of freedom for extension-flexion. The initial angles at these joints were estimated from measured equilibrium leg positions at zero torque. We kept the *CTr* and *TiTa* joints locked in their equilibrium positions. To model the angular spring properties of the leg, we introduce a torsional spring force at each degree-of-freedom with a stiffness that corresponds to each of our experimentally measured values for each leg. For the mesothoracic leg: 8.614 E-9 mN·m/° for levation-depression, 1.01 E-8 mN·m/° for retraction-protraction, 1.202 E-8 mN·m/° for extension-flexion, and 9.873 E-8 mN·m/° for pronation-supination. To simulate the forces arising from the legs contacting with the platform, we introduced Hunt-Crossley forces between the leg tip and platform geometries. We set a maximum simulation time of 0.3 seconds, which we established was enough time for the model to reach a final equilibrium resting position (Figure 4B). We found that the experimental stiffness was not sufficient to support the body, and thus we scaled the stiffness values by different magnitudes to determine how much the stiffness had to be scaled to support the body (Figure 4C). We found that the model transitions to a stable support position – i.e., the model does not fall - when scaling 40-fold. In addition, a scaling value of 1,000 exhibits high stiffness as the equilibrium position approaches the initial equilibrium joint positions. To determine the relationship between the model’s resting equilibrium position and joint stiffness, we plotted scaling values with the model’s final thorax center-of-mass height at the end of 0.3 seconds (Figure 4D).

The fly model with these properties was dropped to the floor. The contact between the ground and each of the legs was described using the Hunt Crossley Forces. The model bounces on the floor a few times before stabilizing at its equilibrium configuration and height.

### 7. OpenSim simulation to determine how fast the active forces decay (Figure 6)

The goal of the OpenSim model in Figure 6 was to assess the time course by which the active forces decay for the median fall. The median height of the fly was 0.5 mm. The height of the fly depends on the active forces before the light is turned on, and the active forces can be predicted from the height of the fly. We assumed that when MNs are inhibited, the active forces will decay exponentially to asymptote to the passive forces. In other words, *f(t)* – the force as a function of time – is given by *f*(*t*) = *f*_*a*_ · *e*^−*t*/*τ*^ + *f*_*p*_. Here *f*_*a*_ and *f*_*p*_ *a*re active and passive forces, respectively. The τ represents the time constant of the decay of the active forces. We simulated the decay with different time-constants ranging from 0.1 seconds to 1 seconds. Data for 0.1 second, 0.2 second, and 0.3 second are presented.

### 8. Electrophysiology recordings of motor neurons

Flies were anesthetized on ice and placed ventral side up in a custom-made chamber. The head was pitched backwards to make room for patch electrode, and the head and thorax was secured in place using wax. The front legs were pulled out and fixed to the side of the body using wax to enable the dissection of the cuticle above the thorax. A small window was made in the cuticle covering the first and second thoracic segment under external saline (103 mmol/L NaCl, 5 mmol/L KCl, 5 mmol/L Tris, 10 mmol/L glucose, 26 mmol/L NaHC03, 1 mmol/L NaH2P04, 1.5 mmol/L CaCl2, 4 mmol/L MgCl2, osmolarity adjusted to 270–285 mOsm, bubbled with 95% O2/5% CO2 to pH 7.1–7.4). The trachea and perineuronal sheath around the first ganglia were removed with fine forceps. Leg motor neurons were targeted for recording by expression of the light-gated anion channel GtACR1 tagged with EYFP, visualized using an Olympus BX51W1 upright microscope equipped with epifluorescence and standard filters. Glass electrodes containing internal solution were used to carry out whole-cell recordings (electrode resistance ranged from 5 to 10 MΩ; 140 mmol/L K-aspartate, 1 mmol/L KCl, 10 mmol/L HEPES, 1 mmol/L EGTA, 0.5 mmol/L Na3GTP, 4 mmol/L MgATP, 265–270 mOsm, pH 7.1–7.4). Voltage was recorded at 10 kHz using a model 2400 patch-clamp amplifier (A-M systems) and low-pass filtered at 5 kHz.

To optogenetically inhibit the motor neuron we used a digital mirror device(DMD) which is a projector that allows the stimulation of any arbitrary ROI. The construction of the digital mirror device has already described in detail recently[32]. During each trial, light spots with 530nm wavelength will be projected onto the patched MN or other neurons in the brain for 5s, then have 5s recovery period and was repeated multiple times.

## Acknowledgment

We would like to acknowledge the members of Bhandawat lab for discussions. Amoolya Tirumalai helped with initial experiments, Kate Medrano with analysis, and Michael Matthews with OpenSim. This research was supported by RO1DC015827 (VB), RO1NS097881 (VB) and an NSF CAREER award (IOS-1652647 to VB).

## Author contributions

VB: Conceptualization; Supervision; Funding acquisition; Writing;analysis. NW: Experimentation, analysis. HB: Experimentation, analysis. JP: Experimentation, analysis;. SBM: Experimentation, analysis; TB: Conceptualization, analysis, writing; AR: Conceptualization, analysis,

## Figure Legends

**Supplementary Video 1. Example video of a tethered fly prep with inactivation phases**. Experimental fly in tethered prep experiences two phases of inactivation with attached weight (0.19mg). Inactivation time is indicated by the “on” cue in the top right corner of the video.

**Supplementary Video 2. Overview of the 3D biomechanical model with simulated joint stiffness parameters**. Video showing an overview of the *Drosophila* 3D biomechanical model developed in OpenSim 4.1. This includes a rotated model of the fly as well as a 100ms simulation of the falling fly with joint stiffness values scaled by 80.

---

**Figure 2-S1.**
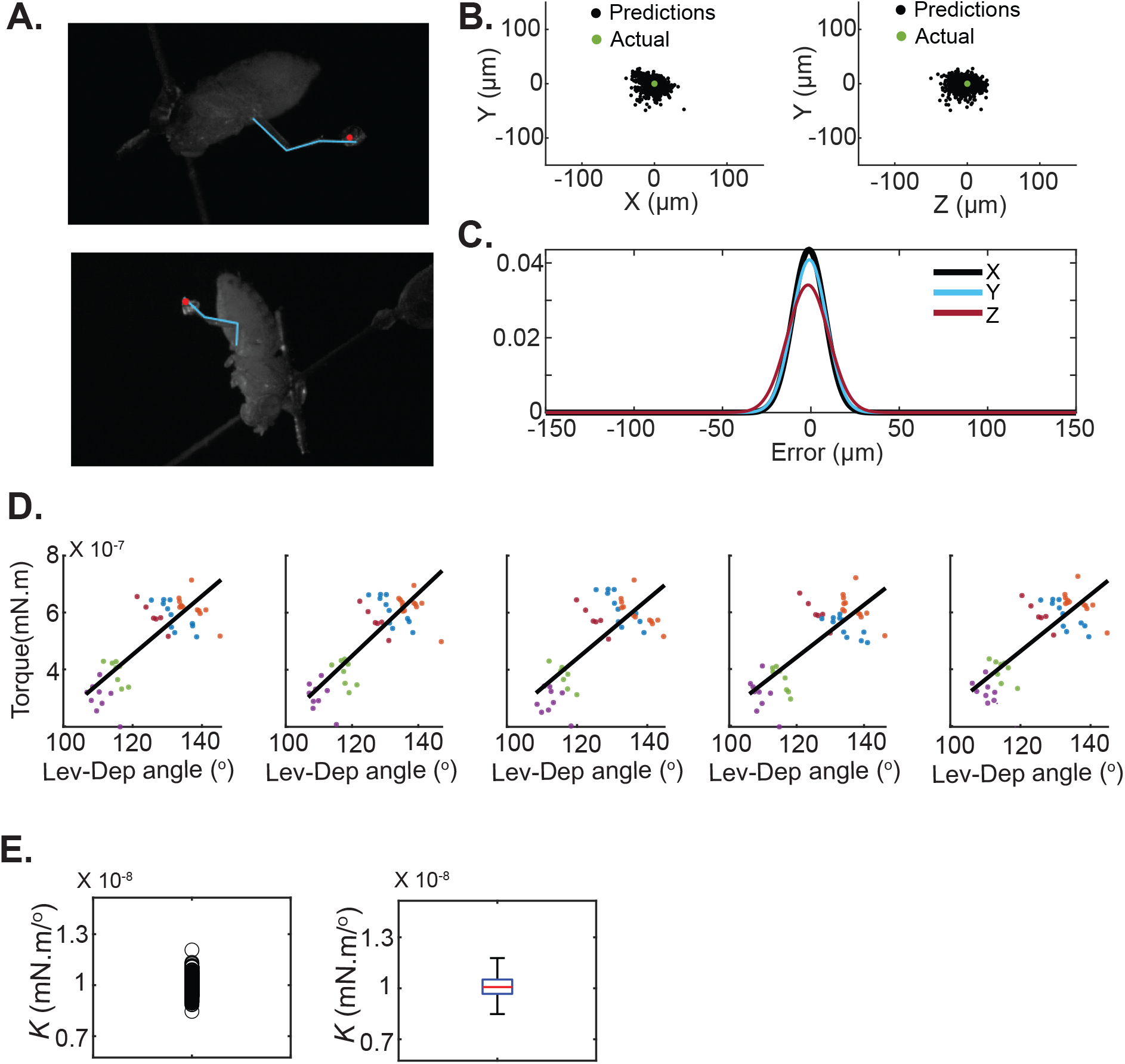
Reconstruction errors resulting from the camera calibration are small. **A**.To test the accuracy of calibration coefficients obtained from the DLT camera calibration, we used inverse DLT to obtain pixel coordinates from the 3D-reconstructed XYZ leg joint positions and overlaid these points on the corresponding frames. The reconstructed leg segments lie on top of the actual leg. **B**.Error distribution across each axis for DLT camera calibrations used for 3D reconstruction of leg kinematics. Error values were obtained by 100 iterations of random subsampling of control points into two sets: The training set for calculating calibration coefficients and the testing set for evaluating prediction accuracy. **C**.The error distribution for each camera calibration is well-captured by a normal distribution. **D**.We sampled from the Gaussian distribution in C to create 1000 simulated legs. This figure shows 4 randomly selected simulations. The first panel shows the original data, while the other four show randomly selected iterations. The variation in results due to DLT error propagation is small. **E**.Final stiffness distributions for levation-depression from all 1000 iterations of DLT error propagation simulations show that variation in results due to DLT error is small.

**Figure 2-S2.**
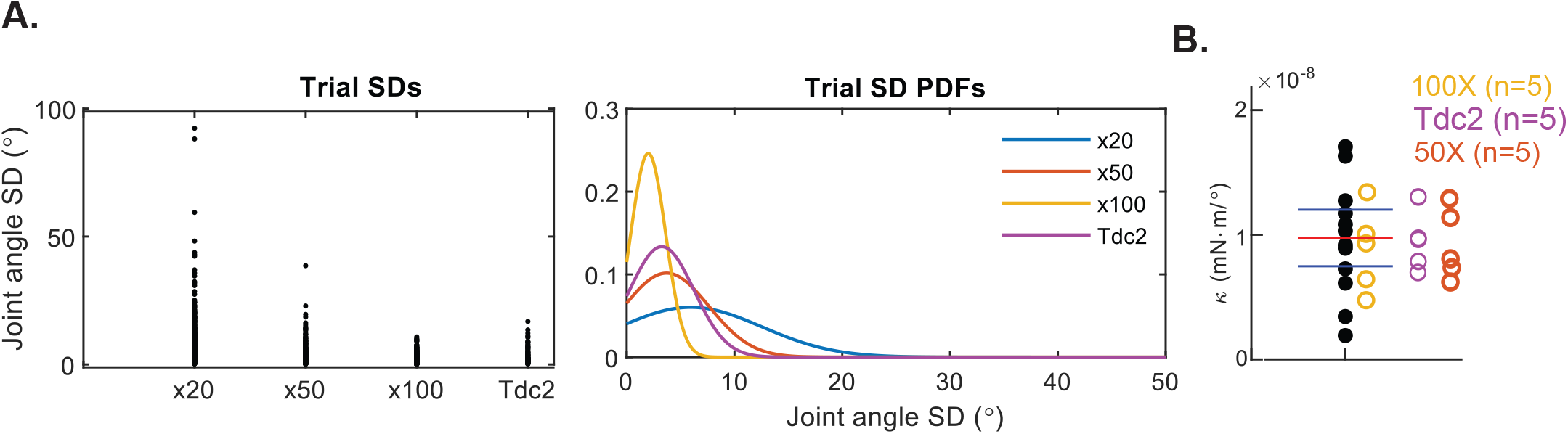
Larger weight and silencing octopaminergic neurons does not affect spring constant measurements. **A**. Standard deviations (SDs) of joint angle values across experimental conditions: x20 etc. refers to the mass of the additional weight. x20, x50, x100 means 20,50,100 times the mass of the leg, respectively. Three masses were used. Tdc2 refers to the case when both glutamatergic and octopaminergic neurons are silenced. Tdc2 experiments were performed with x50 weight. Joint angle SD values per trial for each experimental condition. Probability density functions for trial joint angle SDs by experimental condition. Higher weight reduces the standard deviation. There is no further decrease in SD when both glutamatergic and octopaminergic neurons are silenced. **B**.Stiffness values obtained from experiments with higher weights and when octopaminergic neurons are inactivated. The values lie within the interquartile range observed with the smaller weights. Octopaminergic neuromodulation is silenced optogenetically along with motor neurons using the genetic construct Tdc2-Gal4;UAS-GtACR1(lll) x VGlut(OK371)-Gal4(ll). For all other flies, only motor neurons are optogenetically silenced via the construct VGlut(OK371)-Gal4(ll) x UAS-GtACR1(lll).

**Figure 2-S3.**
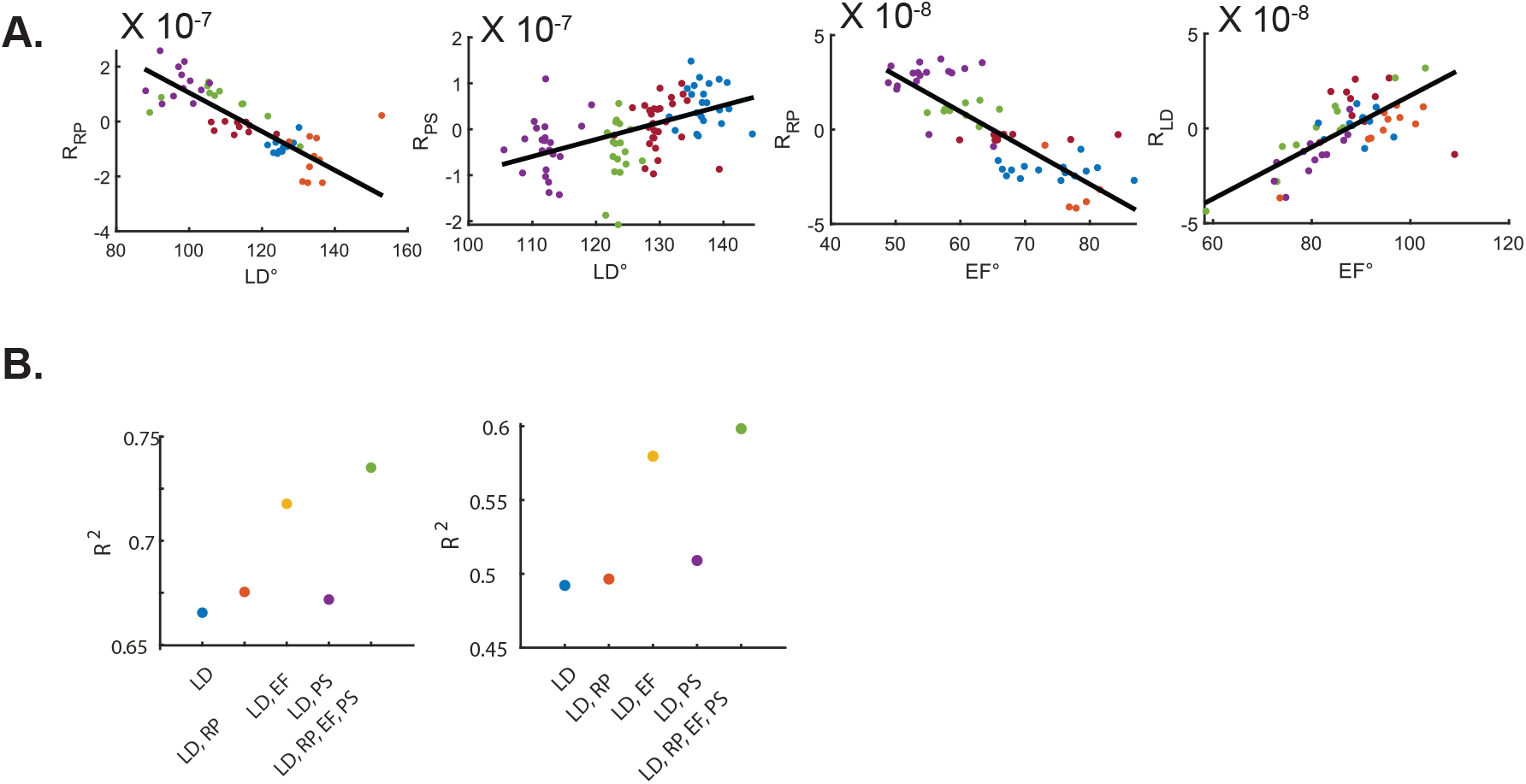
Variation in torque can be explained by the coupled actions of multiple degrees of freedom, as well as other internal forces. **A**.Examples showing that the angle at one degree of freedom is correlated with residuals (torque that is not explained by changes at the original angle) in other degrees of freedom suggesting that torque variation at one degree of freedom (DOF) may depend on other DOFs. As an explicit example of this analysis, for the leftmost panel in A, we first fit the retraction-protraction torque to the changes in retraction-protraction angle as shown in Figures 2-3. Residuals are calculated as the difference between the actual torque and linear fit and are plotted here against the value of the levation-depression angle on the same torque. A significant correlation is found in >60% of cases implying a coupling between the different angles. Importantly, these correlations are likely to be second-order effects. **B**.Using multiple DOF angles to predict levation-depression (LD) torque results in a higher R-squared value than when using only the LD angle. We did not perform this analysis systematically. However, in the 6 flies (just to test the idea), we performed the analysis on we found that there was a much higher correlation when other degrees of freedom were used. Extension-flexion degrees of freedom seem to have a particularly large effect. Levation-depression for two of the flies are shown.

